# The 22q11 low copy repeats are characterized by unprecedented size and structure variability

**DOI:** 10.1101/403873

**Authors:** Wolfram Demaerel, Yulia Mostovoy, Feyza Yilmaz, Lisanne Vervoort, Steven Pastor, Matthew S Hestand, Ann Swillen, Elfi Vergaelen, Elizabeth A. Geiger, Curtis R. Coughlin, Stephen K. Chow, Donna McDonald-McGinn, Bernice Morrow, Pui-Yan Kwok, Ming Xiao, Beverly S. Emanuel, Tamim H. Shaikh, Joris R Vermeesch

## Abstract

Low copy repeats (LCRs) are recognized as a significant source of genomic instability, driving genome variability and evolution. The chromosome 22 LCRs (LCR22s) are amongst the most complex regions in the genome and their structure remains unresolved. These LCR22s mediate non-allelic homologous recombination (NAHR) leading to the 22q11 deletion syndrome (22q11DS), causing the most frequent genomic disorder. Using fiber FISH optical mapping, we have *de novo* assembled the LCR22s in 33 cell lines. We observed a high level of variation in LCR22 structures, including 26 different haplotypes of LCR22A with alleles ranging from 250 Kb to over 2,000 Kb. An additional four haplotypes were detected using Bionano mapping. Further, Bionano maps generated from 154 individuals from different populations suggested significantly different LCR22 haplotype frequencies between populations. Furthermore, haplotype analysis in nine 22q11DS patients resulted in the localization of the NAHR site to a 160 Kb paralog between LCR22A and –D in seven patients and to a 31 Kb region in two individuals with a rearrangement between LCR22A and –B.. This 31 Kb region contains a palindromic AT-rich repeat known to be a driver of chromosomal rearrangements. Our study highlights an unprecedented level of polymorphism in the structure of LCR22s, which are likely still evolving. We present the most comprehensive map of LCR22 variation to date, paving the way towards investigating the role of LCR variation as a driver of 22q11 rearrangements and the phenotypic variability in 22q11DS patients as well as in the general population.

## Introduction

Low copy repeats (LCRs), also referred to as segmental duplications, are a driving force in genome evolution, adaptation, and instability. In the diploid human genome, over 5% of the reference assembly consists of LCRs. This repeat class comprises DNA fragments longer than 1,000 base pairs with ≥90% similarity(*1*–*3*). Duplications have long been recognized as a potential source for the rapid evolution of new genes with novel functions(*3*–*5*). Supporting this dogma, recent studies have indicated potential functional roles for genes within LCRs in synaptogenesis, neuronal migration, and neocortical expansion within the human lineage(*5*–*9*). However, genome assembly builds of these regions are highly enriched for gaps and assembly errors even within the most recent versions of the human reference genome(*10*–*12*). This is because LCRs are both highly sequence identical and copy number polymorphic. These features strongly hamper the study of the precise role of LCRs as drivers of disease or human evolution.

High sequence homology between two (or more) LCR copies is a driver of recurrent genomic rearrangements. Different LCR clusters are often interspersed within individual chromosomes and have originated from several sequential duplication events that occurred during primate evolution(*2*). At loci where paralogous LCRs are on the same chromosome, misalignment of homologous chromosomes or sister chromatids can lead to non-allelic homologous recombination (NAHR)(*13*). Depending on the orientation of LCR paralogs, NAHR between LCRs results in reciprocal deletions, duplications, or inversions. Recurrent deletions and duplications mediated by NAHR have been referred to as genomic disorders(*13*).

The 22q11 deletion syndrome (22q11DS) is the most common genomic disorder, with a prevalence of 1 in 4,000 live births(*14*). The most frequent associated medical features include congenital heart disease, immunodeficiency, palatal anomalies, and hypocalcemia. However, 22q11DS has a heterogeneous presentation and often includes multiple additional congenital anomalies and later onset conditions such as skeletal and renal abnormalities, gastrointestinal differences, and autoimmune disease. In addition, the majority of patients with 22q11DS have developmental delay and cognitive deficits, while a subset have behavioral differences, such as anxiety and attention deficit disorders, as well as, psychiatric illness including schizophrenia in 25%(*14*). The 22q11DS is most often caused by NAHR between LCRs on chromosome 22q11.2 (LCR22s)(*15*). The region contains eight LCR22s, often termed, consecutively, LCR22A-H. In individuals with the 22q11DS, NAHR occurs most often between LCR22A and LCR22D (89%) and between LCR22A and LCR22B in 6% generating a deletion of respectively ∼3 and ∼1.5 Mb. However, other deletions sizes and breakpoints do exist(*16*–*19*). Furthermore, at least three different recurrent translocations between chromosome 22 and chromosomes 8, 11, or 17 have been described in individuals with autoimmune disorders, the Emanuel syndrome, or neurofibromatosis, respectively. The breakpoints of all these translocations are located within palindromic AT-rich repeats (PATRRs), with the one in chromosome 22 localizing to LCR22B(*20*–*24*). PATRRs have been shown to form hairpin or cruciform structures, which are vulnerable to double strand breaks (DSBs), the erroneous repair of which can sometimes lead to chromosomal translocations(*25*).

All LCR22s are a complex of different repeat subunits which are present in variable composition, copy number, and orientation(*26*). LCR22A and LCR22D are the largest and, in the current genome build, estimated to span 1 Mb and 400 Kb, respectively. The size and structure of the LCR22s continues to be variable and inconsistent between various genome assemblies, with the current reference genome still containing sequence gaps within LCR22A. Copy number variations within the LCR22s have been shown to exist by mapping BAC clones derived from different individuals (*27*). In addition, a copy number variant encompassing PRODH, DGCR6, and DGCR5 (referred to as LCR22A+) was recently mapped within LCR22A(*28*). Nonetheless, the overall architecture of several LCR22s remains unresolved. Since the 22q11DS breakpoints are embedded within these unresolved LCR22s, their exact location has, despite extensive efforts (*29*), remained elusive.

To elucidate the LCR22 structures and their variability, and to further refine the 22q11DS rearrangement breakpoint regions, we set out to map these repeats. Using a combination of fiber FISH and Bionano optical mapping we have *de novo* assembled the LCR22s in a diverse set of 186 diploid individuals. Over 30 different LCR22A haplotypes were observed, with alleles ranging in size from 250 Kb to 2,000 Kb, along with the observation of several alternative haplotypes of LCR22D. LCR22A haplotypes varied markedly in their frequencies, with substantial differences between populations. These data have revealed unprecedented variation in LCR size, copy number, and complexity that mediate recurrent genomic rearrangements within 22q11.2. This variation may be a predisposing factor to NAHR and importantly could play a role in modulating the phenotype in both normal and 22q11DS individuals.

## Results

### Subunit-resolution LCR assemblies

To resolve the LCR22 subunit organization, we first redefined repeat subunits that are present in the LCR22s of human genome build 38. By aligning LCR22s and paralogous LCRs to themselves, all 22q11 segmental duplications with a sequence similarity >99% were identified. The LCR decomposition resulted in 24 subunit families composed of sequences with similarity of 99% or higher, 20 of which contain a subunit in the typical 22q11 deletion region (**Table S1**), and 110 low percent identity subunits.

Since the order, copy number, and distances between the subunits in the reference genome is inaccurate, we aimed to resolve the structure by fiber FISH. To visualize the order and distance between those subunits, fluorescent probes were designed for 14 repeat subunits (**Table S2**). These 14 repeat subunits were selected because 1) their size was over 2 Kb, enough to accommodate a visible probe signal, and 2) the subunit copy number was >1 and < 10 on chromosome 22q11.21 (using BLAT), so that probes were visually countable and evenly spread over the LCR22s. A genome wide hg38 BLAT alignment of those 14 subunit probe sequences generated 82 paralogous hits larger than 1 Kb, of which 67 were specific for the LCR22s. Probes were generated by long range PCR, capturing individual subunit families and subsequently labeled. To guarantee correct identification of signals, different subunits that flank each other in LCR22A (hg38) were designed to have different colors (**Fig. 1C, UCSC track)**. BAC probes distal and proximal to the LCR22 clusters were used to anchor any detected repeat patterns to unique sequence.

**Fig. 1.**
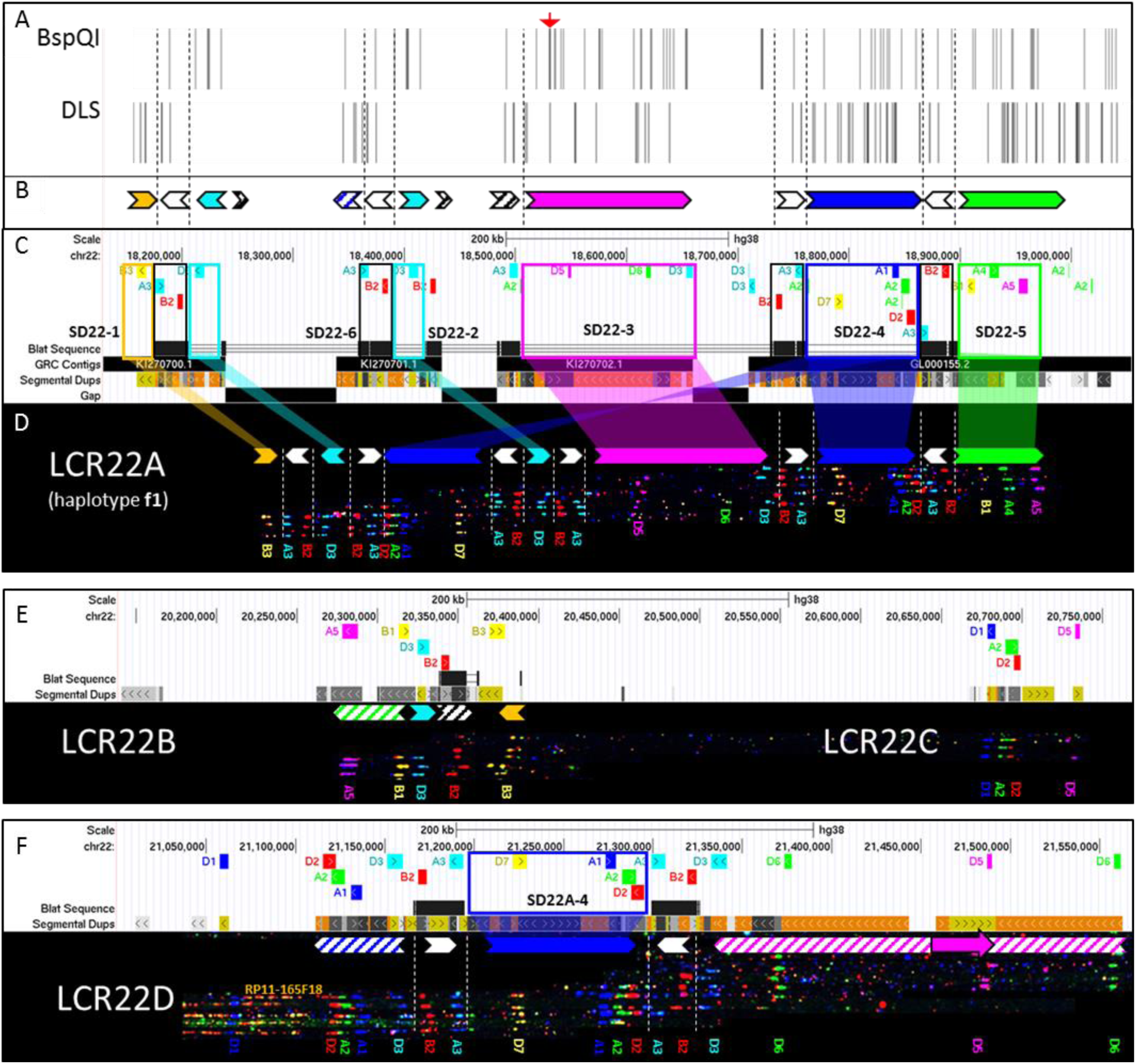
*In silico* hg38 fiber FISH probe positions compared to duplicon composition of LCR22A. A) Refseq curated gene set overlapping with the LCR22s. B) Duplicon decomposition of the hg38 structure of LCR22A. Duplicons were deduced from mapped haplotypes. Filled, colored arrows represent copies of duplicons and hatched arrows represent partial copies of duplicons of the same color. C) UCSC browser hg38 reference assembly tracks of Segmental Dups^2,24^, GRC contigs, gap positions, and fiber FISH probe BLAT positions (white panel). Positions of the latter are aligned with recordings of fiber FISH patterns in LCR22A (black bar). Decomposition of one LCR22A haplotype to duplicons is illustrated using colored (non-white) arrows. For the showcased allele, duplicon order centromeric to telomeric is SD22-1, -2(-), -4(-), -2(+), -3, -4(+), and -5, where ‘-‘ represents inverted and ‘+’ represents direct orientation. Larger duplicons are flanked by copies of SD22-6 (white arrows). Probe identifiers are indicated below the fiber pattern. D) Refseq annotated genes overlapping with LCR22B and –C. E) LCR22B and – C, fiber FISH patterns have the same order and distances as those predicted in hg38 and contain partial duplication of LCR22A duplicons (hatched arrows). F) Refseq annotated genes overlapping with LCR22D. G) all LCR22D molecules present at the same centromeric start, overlapping with predicted hg38 probe positions. The first duplicon displays a partial SD22-4 and SD22-2 (hatched blue arrow), followed by a complete SD22-4 flanked by SD22-6 copies (white arrow). The distal end of LCR22D consists of partial duplications of SD22-3 (hatched magenta arrow). Nested, solid magenta arrow represents probe D5 position variant.

The LCR22-specific fiber FISH probe pattern was first assayed on DNA fibers generated from haploid cell line CHM1 and HapMap cell line GM12878. These genomes have been well characterized and were included in the Platinum Genome Project(*30*). The haploid state of CHM1 significantly reduced mapping complexity of the repeat clusters. Hybridization of our custom probe set on fibers of CHM1 and GM12878 resulted in over 100 informative fibers, each over 200 Kb in size. By overlapping patterns identical in color sequence and spacing, chimeric fiber patterns and false positive signals caused by noise, were eliminated. Tiling the clustered fibers enabled *de novo* assembly of the subunit order over more than 1 Mb.

To link the LCR22 repeats to unique sequence, BACs flanking the repeats were used as probes and allowed identification of the assembled LCR22s (**Fig. S1).** Assembled subunit patterns were compared to the *in silico* defined subunit positions in the reference genome. Probe patterns of LCR22B and LCR22C were in agreement with those in hg38 for both cell lines (**Fig. 1E**). In contrast, LCR22A diverged from the (incomplete) reference genome, with some divergence also in LCR22D (**Fig. S2**, **Fig. S3**, **and Fig. S4**). LCR22D structure was identical between both cell lines, with the position of the proximal fifteen fluorescent signals and the most distal signal matching those present in the reference genome. A single LCR22D probe, D5, presented with a different position in all assembled alleles from these two cell lines (**Fig. S2B**). In LCR22A, the d*e novo* assembled probe pattern in CHM1 contained 47 probe signals, as opposed to the 23 signals predicted in hg38 (**Fig. S3**). Of those 47, signal colors and spacing matched hg38 for the first 4 and the last 12 probes. GM12878 presented two alleles, consisting of 47 fluorescent signals on one allele and 30 on the other (**Fig. S4**). On the first allele of LCR22A, 14 red, 6 green, 4 blue, 15 cyan, 2 magenta, and 6 yellow signals were counted. However, the order of, and distance between-, signals only matched the reference proximally for the first 4 proximal and the last 12 distal probe signals in the LCR. The second allele of LCR22A produced 9 red, 4 green, 3 blue, 9 cyan, 1 magenta, and 4 yellow signals, overall fewer than the first. Similar to the first allele, the first 4 proximal and 12 distal fluorescent signals matched with the reference genome, while the signals in between differed from the positions represented in hg38. Based on the distance between probe signals that matched the reference, the total LCR22A sizes in GM12878 could be estimated as ∼1.20 Mb for the first and ∼0.65 Mb for the second allele. The CHM1 allele had a length around 1.20 Mb as well, although its composition differed from the GM12878 alleles.

Since those first three LCR22A assemblies differed substantially from the reference and from one another, we wondered whether those alleles were exceptional. To evaluate variation more broadly, we assembled fiber patterns in 29 cell lines, resulting in an additional 41 assembled molecules. For LCR22B and LCR22C patterns were identical in all samples. In contrast, we observed three haplotypes for LCR22D (**Fig. 2B**) and a staggering number of haplotype variants for LCR22A (**Fig. 2A**). The four most proximal and five most distal probe signals of LCR22A always matched the reference positions. In contrast, the remainder of LCR22A alleles showed astonishing variation: In 44 LCR22A alleles, 26 distinct haplotypes were observed, varying in length from ∼300 Kb to over 2 Mb (**Fig. 2**, **Fig. S5**). No individual in this cohort was homozygous for the structure of LCR22A. In contrast, LCR22D displayed less variability, with allelic variation including different positions of probe D5, and a duplication of probes A1-A2-D2-A3-B2-D3 (**Fig. S2C)**.

**Fig. 2.**
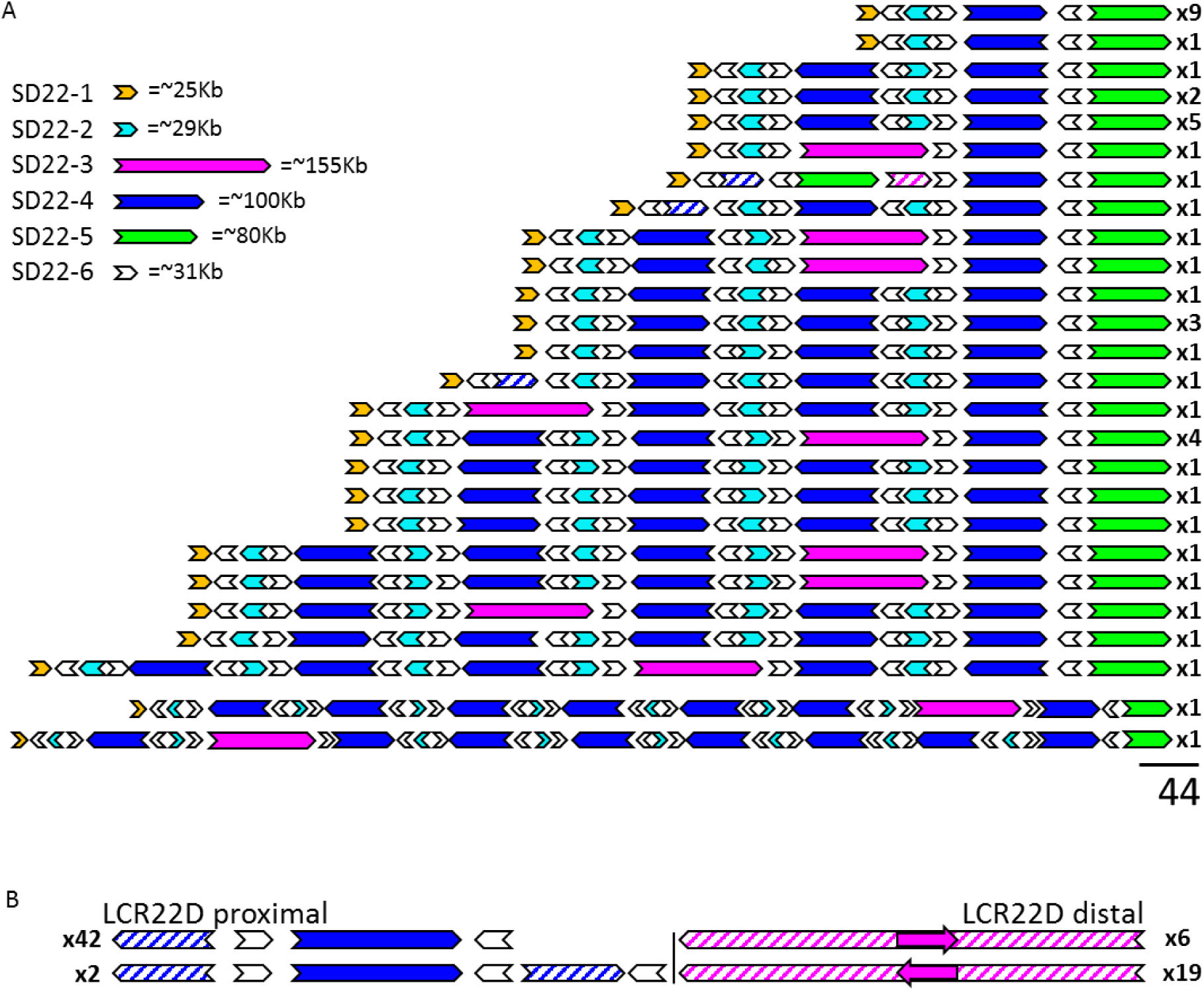
Fiber FISH mapped haplotypes of LCR22A and LCR22D observed in a cohort of 19 diploid individuals, 1 molar cell line, and 5 non-rearranged alleles from22q11DS probands. A) 26 haplotypes observed for LCR22A. Haplotypes are aligned at the distal unique anchor of LCR22A. B) Proximal and distal haplotypes observed for LCR22D. Filled, colored arrows represent copies of duplicons, hatched arrows represent partial copies of duplicons of the same color. Size estimates of individual SD22s are shown (upper left). Frequencies of haplotypes are depicted on the right (i.e. x9, x1, etc.).

### LCR22A fiber patterns identify core duplicons

Despite the observed scale of variation, mapped haplotypes displayed clusters of conserved, non-random probe patterns. Copy number variation of a small set of duplicons could drive the observed differences in LCR22 architecture. To uncover this core set, we visually deduced a minimal set of different probe clusters that could explain the composition of a maximal number of LCR22 haplotypes. Those probe clusters represent segmental duplications of the LCR22s, so we designated them as SD22s.

We identified six probe clusters to define six LCR22 duplicons, designated SD22-1 to -6. All 44 fiber FISH mapped alleles of LCR22A presented a conserved proximal and distal end. Yellow probe B3 (SD22-1) marked the centromeric unique anchor, while a yellow-green-magenta pattern (B1, A4, A5; SD22-5) was observed at the telomeric unique anchor (**Fig. 1**). Between those anchors, four probe clusters were respectively characterized by a cyan signal between two red probes (D3; SD22-2), a magenta-green-cyan cluster (D5, D6, D3; SD22-3), a yellow-blue-green-red cluster (D7, A1, A2, and D2; SD22-4), and a cyan-red cluster (A3 and B2; SD22-6) (**Fig. 1B,C**). In contrast to the proximal and distal anchors, those four probe clusters were copy number variable amongst alleles, with SD22-3 being absent in some (**Fig. 2A**). However, their individual structural integrity, was retained when comparing different LCR22 alleles, and these clusters were only occasionally partially represented. In three LCR22A haplotypes, a partial duplication of SD22-4 and -2 was present. This fragment only contained probes A1, A2, and D2 of the SD22-4 probe cluster and D3 of the SD22-2 cluster.

This analysis revealed that every LCR22A allele mapped by fiber-FISH was a composition of SD22-1 to -6 in different orders, copy numbers and orientations. Notably, the order of SD22s was non-random. No tandem repeats of single duplicons were observed. Instead, any two subsequent copies of SD22-1, -2, -3, -4, or -5 were always flanked by a paralog of SD22-6. Moreover, the orientation of SD22-6 relative to its surrounding duplicons was conserved. Flanking SD22-4, both SD22-6 paralogs always faced inwards, SD22-2 was flanked by two SD22-6 paralogs facing outwards, and SD22-3 had both flanking SD22-6 paralogs in a direct (+) orientation. The unique anchor SD22-1 was followed by an inverted SD22-6 and likewise, SD22-5 was preceded by an inverted (-) paralog of SD22-6. Although most molecules present a single copy of SD22-1 and -5 at respectively the centromeric and telomeric end of assembled LCR22A haplotypes, copy number variants of SD22-5 do exist as well. Previously, a deletion (0.3%) and reciprocal duplication (1.3%) embedded in LCR22A were identified in a cohort of 15,579 normal individuals(*28*). An individual that was previously determined to carry such a duplication polymorphism was included in the mapped cohort (**Fig. S9, family 5, A**). A fiber FISH map of this individual’s LCR22A confirmed a SD22-5 duplication within LCR22A. The duplicated fragment included the probe pattern of SD22-5, in which PRODH, DGCR5, and DGCR6 are embedded. Rather than being duplicated in tandem, both copies were separated by a fragment of SD22-3 and a copy of SD22-4.

By comparing the probe patterns to matching positions in the reference genome, the associated structure based on the reference genome sequence was deduced. *In silico* fiber FISH probe patterns of reference contigs KI270701.1, KI270702.1, and the proximal 150 Kb of reference contig GL000155.2 individually matched duplicons SD22-2, -3, and -4 respectively (**Fig. 1B,C)**. Flanking those duplicons, SD22-6 corresponded to a ∼31 Kb repeat in hg38, which was present five times in the reference LCR22A with sequence similarities of 97% and higher (**Fig. 1B,C, blat track and white arrows)**. SD22-6 flanked both sides of SD22-4 in opposite orientations, and also flanked the internal sides of centromeric and telomeric anchors SD22-1 and SD22-5. The SD22-6 hg38 sequence contains paralogs of a lincRNA with sequence similarity to FAM230C. Each of these paralogs contains segmental duplications of the translocation breakpoint type A (TBTA, AB261997.1), which is constituted of an unstable palindromic AT-rich repeat (PATRR).

### Bionano analysis confirms the fiberFISH *de novo* assemblies

To evaluate the fiberFISH assemblies with an orthogonal technology, Bionano assays were performed on the haploid cell line CHM1, GM12878, and two parent-22q11DS child trios. The fiber FISH and Bionano results were compared by converting the fiber FISH duplicon order and orientation information into sequences, stitching together the duplicon sequences from the reference genome. These sequences were *in silico* labeled to convert them to Bionano optical map format (**Fig. S6B, E, H, L, O, R, U, X, BB, EE, HH, KK, green bar)** and then compared to the observed Bionano assemblies in the same individual (**Fig. S6b, E, H, L, O, R, U, X, BB, EE, HH, KK, blue bar)**. Since a certain degree of paralogous variation between segmental duplications is missed at the fiberFISH resolution, some mismatch is expected when comparing individual label sites(*2*). However, outside of this expected paralogous variation, *de novo* assembled LCR22 duplicon order and orientation should be consistent in both datasets.

For CHM1, GM12878, and the two trios, Bionano data were generated using the DLS (“direct label and stain”) enzyme. After examination of single molecules from these samples at both LCR22A and LCR22D, a list of partial haplotypes with strong single molecule support was generated for each sample (**Fig. S6**). Next, we noted that the cluster of SD22-4 and its two flanking SD22-6 paralogs together revealed five DLS labels that were polymorphic between paralogs (**Fig. S7**). These polymorphisms were then used to stitch the partial haplotypes into end-to-end haplotypes of LCR22A and LCR22D.

Observed optical maps from CHM1, GM12878, and all 12 LCR22A alleles in the two families were aligned to those converted from the fiber FISH results (**Fig. S3**, **Fig. S4**, **Fig. S6**). All of those alignments showed strong agreement between *in silico* and observed optical maps after accounting for the expected paralogous variation. The copy number, order, and orientation of detected duplicons was identical between the two techniques. As expected, discrepancies typically presented as single labels that were detected in only one of the two datasets, and occurred on average six times every megabase (81 discrepant sites/ 12.42 Mb of LCR22A assemblies). Most of these discrepant positions overlapped with polymorphic sites detected with the DLS marker, which are below the resolution of the fiber FISH assay (**Fig. S7**). A consistent size discrepancy between converted and observed maps was also present in SD22-3.

### Bionano optical mapping reveals population specific structural variation at LCR22A

To determine the prevalence of different variants in LCR22-A and –D, both within and between populations, the variability was mapped by Bionano optical mapping using the Nt.BspQI enzyme, in a cohort of 154 phenotypically normal individuals from 26 populations spanning five super-populations: African (AFR), American (AMR), East Asian (EAS), European (EUR) and South Asian (SAS) (Levy-Sakin *et al*. 2018). Optical map assembled contigs that overlapped LCR22A and had single molecule support, were collapsed into distinct configurations (**Fig. 3A-D).** Not all assembled contigs were able to span the entire LCR22A region from end to end, due to factors such as insufficient molecular coverage, short molecule lengths, and/or longer LCR22A haplotype lengths. Additionally, the presence of two Nt.BspQI nicking sites in close proximity to one another near the beginning of SD22-3 created a fragile site on the DNA that consistently interrupted molecules in that location **(Fig. S8A)**. While this catalog of LCR22A haplotypes is likely not comprehensive, it captures a substantial amount of large-scale structural variation at this locus.

**Fig. 3.**
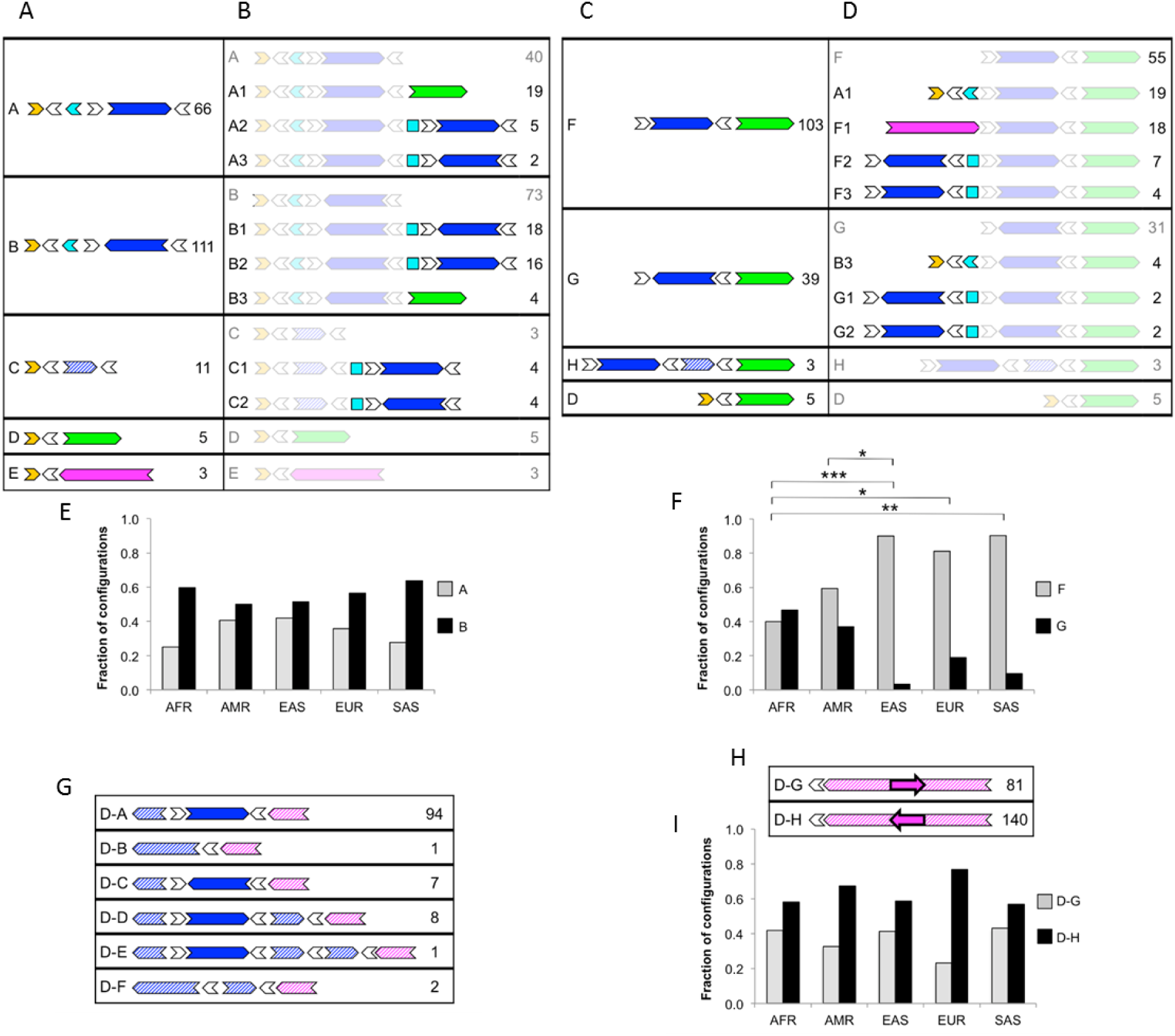
LCR22A and LCR22D configurations across a diverse control dataset observed using Bionano optical mapping. Diagrams depict order and orientation of observed duplicons as defined in Fig. 1. A-B) Minimal (A) and extended (B) configurations anchored in unique sequences proximal to LCR22A. C-D) Minimal (C) and extended (D) configurations anchored in unique sequence distal to LCR22A. E) Population distribution of occurrences for proximal LCR22A configurations A and B. F) Population distribution of occurrences for distal LCR22A configurations F and G. G) Configurations anchored in unique sequence proximal to LCR22D. H) Configurations anchored in unique sequence distal to LCR22D. I) Population distribution of occurrences for distal LCR22D configurations D-G and D-H. For each configuration in a-d and g-h, the ID is on the left and number of times that configuration was observed in the dataset is on the right. Duplicons for which an orientation could not be determined are represented as squares. Population abbreviations: AFR, African; AMR, American; EAS, East Asian; EUR, European; SAS, South Asian. * *p* < 0.05, ** *p* < 0.01, *** *p* < 0.001, Fisher’s exact test, adjusted.

A total of 16 non-redundant partial and complete LCR22A configurations were identified in the set of 154 individuals (**Fig. 3**). As seen in the fiber FISH results, the majority of the variation involved variable copy number and orientation of SD22-4. The predominant variants began with the duplicons SD22-1 and -2, followed by SD22-4 in either direct or inverted orientation, with copies of SD22-6 flanking each duplicon (**Fig. 3, ‘A’ and ‘B’ configurations)**. Consistent with the fiber-FISH alleles, other configurations followed the anchoring SD22-1 duplicon with either a partial copy of SD22-4 (configuration C) or a copy of SD22-3 (configuration E). One notable configuration, not observed in the fiber FISH dataset, harbored a deletion of almost the entire locus, with a minimal composition of SD22-1 to inverted SD22-6 to SD22-5 (configuration D). Configurations anchored downstream of LCR22A (**Fig. 3 c-d)**, showed similar patterns, preceding the anchoring SD22-5 with either a full or partial copy of SD22-4 in direct or inverted orientation (configurations F-H). All repeat duplicons were flanked by copies of SD22-6.

After compiling this set of distinct configurations, we next set out to evaluate their prevalence within the dataset. Since assembled optical map contigs were prone to assembly errors at long segmental duplications, we instead analyzed single molecules from each sample. In order to separate partial configurations from longer ones of which they are subsets, the configurations were divided into groups, first into those anchored in unique sequences proximal or distal to LCR22A and then into minimal (**Fig. 3A,C**) and extended (**Fig. 3B,D**) configurations, with some end-to-end haplotypes falling into both proximal and distal groups (e.g. configurations A1 and B3 in Fig. 3 b and d). Local single molecules from each sample were then aligned to all of the possible configurations within the group and the best match was selected (see Methods for additional details) in order to determine the configuration(s) present in each sample.

The results showed dramatic differences in prevalence between different configurations (**Fig. 3**). For minimal configurations at the proximal end of LCR22A (**Fig. 3A)**, the most common duplicon to follow the initial cluster of SD22-1, SD22-2, and flanking SD22-6 copies was an inverted copy of SD22-4 (configuration B in **Fig. 3A**), which accounted for 111/196 (57%) and 26/44 (59%) of the configurations in this group, in optical map and fiber FISH datasets respectively. Next most common at approximately half the frequency (34% in optical map data and 31% in fiberFISH data), was a copy of SD22-4 in the direct orientation (configuration A in **Fig. 3A**), i.e. the structure corresponding to the beginning of both the hg19 and hg38 reference haplotypes. These results indicated that neither of the two most recent reference genomes represented the major allele at this locus. Among the minimal configurations anchored downstream of LCR22A (**Fig. 3C)**, a direct copy of SD22-4 preceded SD22-5 in 103/150 cases (69%, configuration F), a configuration that is consistent with the reference genomes. The next most common configuration (configuration G, 26%) had SD22-5 preceded by an inverted SD22-4. These frequencies differed significantly (*p* = 0.041, Fisher’s exact test) in the fiber FISH dataset, in which 39/44 (89%) of chromosomes display configuration F while only 5/44 (11%) showed SD22-5 preceded by an inverted SD22-4. Notably, the fiber FISH samples were taken exclusively from individuals of European descent, and the distal portion of LCR22-A differed significantly among various ethnicities (**Fig. 3F**). Among European samples in the Bionano dataset, configuration F was seen in 79% of alleles (26/33 cases) while G was seen in the remaining 21% (7/33 cases), showing no significant difference compared to the fiber FISH results (*p* = 0.34, Fisher’s exact test). For both groups of minimal configurations, the shortest end-to-end haplotype containing SD22-1 to inverted SD22-6 to SD22-5 (configuration D) comprised a small minority of cases, with a prevalence of under 4%.

The extended configurations were able to be observed in fewer samples than the minimal ones because not all single molecules that matched the minimal configurations were long enough to extend into an additional duplicon. Nevertheless, this smaller dataset illustrated several distinctive patterns. The three most common proximal configurations, each representing ∼20-24% of the extended proximal alleles, followed the anchoring SD22-1 and SD22-2 with: 1) direct SD22-4 and SD22-5, i.e. the hg19 haplotype (**Fig. 3B**, configuration A1); 2) tandem copies of indirect SD22-4 (configuration B1), and 3) an indirect and then a direct copy of SD22-4 (configuration B2). The remaining 34% of cases comprised seven configurations, each with 3-6% prevalence. Among the distal extended configurations (**Fig. 3D),** the two predominant configurations, each with 28-30% prevalence, were the hg19 haplotype (configuration A1) and a configuration (F1) matching the distal end of the hg38 haplotype, i.e. SD22-3, SD22-4, and SD22-5. Although the latter configuration was also common in the fiber FISH data (23% of alleles, **Fig. 2A**), the ten end-to-end haplotypes containing configuration F1 showed only one match to the exact hg38 structure, instead representing a total of seven haplotypes of varying lengths and copy number of SD22-4, suggesting that this configuration class in the optical map data is likely to also represent a wide range of end-to-end haplotypes.

Having determined configuration frequencies in the entire dataset, we next wanted to determine whether configuration frequencies differed by population. To increase sample size and power, we focused on the two most common minimal configurations at either the proximal or distal end of LCR22A, i.e. those involving complete copies of SD22-4 in either orientation (configurations A, B, F, and G). The results showed substantial differences between super-populations for the distal configurations (*p* < 0.05, Fisher’s exact test), with the largest difference occurring between the African and East Asian populations (**Fig. 3F)**. Thus, at the distal end, while SD22-4 in the direct orientation was more common overall, it represented only 42% (11/26) of African haplotypes, compared to 85% (22/26) of East Asian haplotypes (**Fig. 3F**). At the proximal end, SD22-4 in an inverted orientation was more common than in direct orientation in every population (**Fig. 3E)**.

### Structural variation at 22q11 LCR22D

LCR22D was substantially less polymorphic and complex than LCR22A, but it nonetheless harbored some large-scale structural variation. Following the same procedure as above, haplotypes for LCR22D were compiled from the optical map data from 154 individuals. Six proximally-anchored configurations were observed (D-A to –F), five of which involved the paralog of SD22-4 that is present in the proximal half of LCR22D (**Fig. 3G)**. In the distally-anchored half of this locus, only a 64 Kb inversion was observed (**Fig. 3H).** Since these two regions were distant from one another, they were analyzed separately in order to minimize the length of the molecules required to identify each configuration. In the proximally-anchored half, the reference haplotype (D-A) was predominant, accounting for 83% of all alleles (compared to 95%, 42/44, in the fiber FISH data **Fig. 2B**). Among the other configurations, a full inversion or partial duplication of SD22-4 (D-C and D-D) were the most common, with frequencies of 6% and 7%, respectively **(Fig. 3G)**. In the fiber FISH data, a partial duplication (D-D) is observed in 2/44 (4.5%) alleles, however, no inversions of SD22-4 were observed. We detected no population-based differences among these proximal configurations.

In the distally-anchored half of LCR22D, the inversion configuration (D-H) was more common than the reference (D-G), with an overall prevalence of 63% (140/221). The inversion prevalence in European samples was 77% (30/39), consistent with its prevalence in the European samples mapped by fiber FISH (76%, 19/25 cases). In the optical map cohort, the D-H variant was the major allele in all five super-populations, with significant differences between super-populations observed in the dataset as a whole (*p* < 0.05, Fisher’s exact test), although pairwise differences between super-populations did not reach statistical significance after multiple testing correction **(Fig. 3I)**.

### The 22q11DS rearrangement breakpoints are localized within SD22-4 and flanking SD22-6

Thus far, it remained impossible to map the 22q11DS rearrangement breakpoints within the LCR22s. To resolve those rearranged alleles, refine the breakpoint region, and map potential variability of the rearrangements, we generated assemblies using either a combination of fiber FISH and optical mapping or fiber FISH only for 22q11DS patients from eight families (**Fig. S9**). To reduce complexity and assure correct assembly of the rearranged LCR22s, fiber FISH maps were generated from somatic cell hybrids containing only the del(22)(q11.21), derived from lymphocytes of the probands in Families 1 and 2. For both of these probands, patterns of the rearranged LCR22s were reflected identically in the somatic hybrids and EBV cell lines (**Fig. S10**). In all families, the parent of origin of the deletion was also confirmed by STR marker analysis (data not shown).

Seven of the probands tested had the typical LCR22A-D deletion, while one proband carried the smaller LCR22A-B rearrangement (**Fig. 4, Fig. S9**). In Family 1, the individual with the 22q11DS presented with five LCR22 patterns. Four were indicative of complete alleles of LCR22A, -B, -C, and -D. Moreover, these were all identical to one of the mother’s LCR22 structures, and thus represented the non-rearranged allele (**Fig. 4 B,D**, **Fig. S9 B,D)**. The fifth LCR22 pattern initially presented with a duplicon order and orientation identical to one of the father’s LCR22A alleles. Distally from the third copy of SD22-4, the allele transitioned to a pattern identical to one of the LCR22D alleles in the same parent. The probe pattern suggested that LCR22A and –D had been merged into one LCR, with a breakpoint either in SD22-4 or in one of its flanking SD22-6 paralogs.

**Fig. 4.**
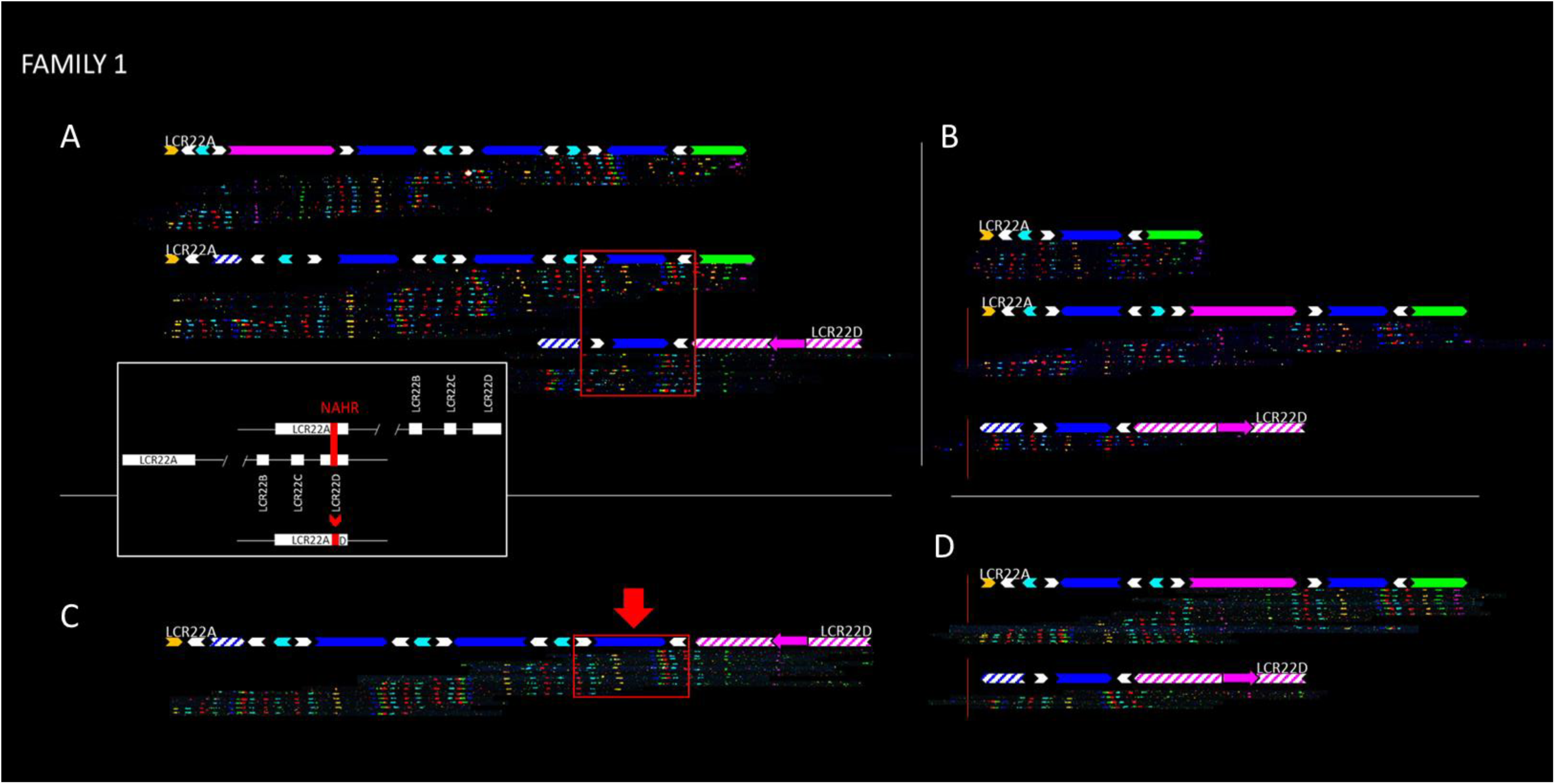
Analysis of a 22q11.21 deletion in a proband and parents using fiber FISH. The family shown here is family 1 in Fig. S9. Fiber FISH assemblies are aligned with a duplicon representation as defined in Fig. 1. Both alleles of LCR22A and LCR22D are shown for each individual in the family trio. All individuals had the same configuration at both alleles of LCR22D which is thus shown only once for each. A) Parent of origin. The shared region between LCR22A and –D is marked by a red box. B) Other parent. Non-rearranged alleles of LCR22A and –D transmitted to proband are shown inside red box. C) Rearranged and D) non-rearranged alleles observed in proband. Red arrow marks duplicon(s) that were involved in recombination and define the breakpoint region. The white inset depicts a schematic overview of NAHR between LCR22A and –D.

Within the probands, seven out of eight of the rearranged alleles overlapped with the parental alleles at the longest shared region between LCR22A and -D (**Fig. 1**, **Fig. 4**, **Fig. S9**), suggesting this as the likely location for the rearrangement breakpoints. This region comprises SD22-4 and two flanking SD22-6 duplicons, together forming a 160 Kb stretch of homologous sequence. These constitute a large direct repeat in LCR22A and –D, which has been previously proposed to contain rearrangement breakpoints(*29*). In all individuals, one copy of SD22-4 was present in LCR22D in a direct orientation (**Fig. S9**), while it is found in variable copy numbers and orientations in LCR22A. In all seven rearranged alleles, SD22-5 was deleted, thereby generating hemizygosity for PRODH, DGCR5, and DGCR6. The LCR22A-B rearrangement was mapped in one family (**Family 7, Fig. S9**). One of the father’s LCR22A alleles was identical to his offspring’s rearranged chromosome, up to the last distal direct paralog of SD22-6 (white arrow). At this position, the patient’s rearranged LCR22 transitioned to the last two probes of LCR22B. This pattern suggested that a non-allelic homologous recombination occurred at SD22-6. Assembly of a second individual with an LCR22A-B deletion supports this breakpoint location distally in the fragment of duplicon SD22-6 (**Fig. S11).**

### Discussion

The LCR22 reference sequences have contained gaps since the first human genome assembly was released. We now show that an astounding level of inter-individual variability of LCR22A, and to a lesser extent LCR22D, has likely impeded the assembly of a complete reference sequence for these LCRs. Further, although whole-genome short-read sequencing is now routine, alignment of short sequencing reads to the human reference sequence generally fails to detect and assemble large structural variants and repetitive regions like the LCR22s. Due to the length of the duplications, even assemblies using longer-range technologies like PacBio and 10X Genomics linked reads have been unable to assemble these regions(*31*, *32*). To resolve these problems, we have generated a *de novo* assembly of ? complete LCR22 haplotypes and partial LCR22 configurations in 154 diverse diploid individuals by combining fiber FISH and Bionano optical mapping. These maps revealed at least 30 different alleles of LCR22A and six variants of LCR22D. LCR22A alleles observed in our study ranged in size from ∼250 to ∼2000 Kb. Most of these alleles could be decomposed into six core duplicons (SD22-1 to -6), with duplicons presenting in different orientations and at variable positions within the LCR. The most frequent LCR22A haplotype had the following structure; SD22-1, -6(-), -2(-), -6(+), -4(+), -6(-), -5, which made up ∼ 25% of all mapped alleles. Notably, its structure is remarkably similar to one of the first LCR22A sequences proposed in Shaikh *et al.* 2000(*17*), a haplotype which was presented in hg19. Thus far, only one smaller haplotype was detected, in which SD22-1 was directly followed by SD22-6(-) and -5. This might indicate the requirement of a minimal haplotype to maintain a viable gene dosage.

Since our sample size is relatively small, we expect that the alleles we observed are likely to be a small subset of all possible haplotypes that may exist in the population. The fact that none of the 24 individuals tested were homozygous for LCR22-A, further supports the existence a high number of different haplotypes in humans. Consequently, any homologous recombination between two (different or identical) alleles of LCR22A will likely generate a novel allele with a duplicon composition different from the parent of origin (**Fig. S12**). Notably, LCR22s are known to be sites with an increased recombination rate when compared to their surrounding loci(*33*, *34*). Thus, LCR22s are likely to be ‘hot-spots’ for the introduction of novel structural variants in the population. Furthermore, the various configurations of LCR22A and -D varied in frequency amongst populations. It is likely that some of these configurations are more vulnerable to NAHR than others. Therefore, it is possible that the variation in the frequency of the 22q11DS amongst populations(*35*, *36*), may result from the differences in the frequencies of various LCR22A and -D configurations and their respective vulnerability to NAHR.

Studies on genome-wide LCR diversity have identified numerous LCR clusters, mainly in pericentromeric and subtelomeric regions(*37*). However, none of those come close to the level of complexity and the unprecedented number of haplotypes found in the LCR22s. A few studies using either WGS read depth based predictions, digital droplet PCR, or custom BAC arrays have revealed copy number variability between individuals within regions containing LCRs(*38*–*41*). Eight distinct haplotypes have been described for the LCR clusters on chromosome 17q21.31, ranging in size from 1.08 to 1.49 Mb(*42*). Similarly, copy number variation, ranging from 2-11 copies of a ∼900-Kbp region (chr15:20353991-27802370) in 15q11-q12 was observed in the 1000 genome project(*43*). Individuals at the extremes of this spectrum, i.e 2 copies versus 11 copies, are estimated to differ by ∼8.1 Mb of DNA(*44*). Such repeat expansions have mainly been found to be human-specific when compared to their orthologs in great ape genomes (*45*). Moreover, significant variation between different human populations suggests that these genomic rearrangements happened recently or are still ongoing(*40*). However, a majority of these studies are based on short read whole genome sequencing data, which are not as reliable for determining true copy number and complex architecture of regions containing LCRs.

Our approach has also allowed us to further refine the localization of recurrent rearrangements to specific modules within LCR22. In seven out of eight families, the rearrangement occurred within a 160 Kb core module, containing SD22-4 and SD22-6, which is present within both LCR22A and -D in all different haplotypes. A previous study had predicted that NAHR occurred within a duplicon referred to as BCRP2 (**Family 7 and 8; Fig. S9)**(*29*). Our analysis of the same two trios confirmed the presence of paralogous repeats of BCRP2 at several positions in the LCR22s and further showed that NAHR occurred between one predicted and one BCRP2 locus embedded in the SD22-4 duplicon. In two families with LCR22A-B deletions the breakpoint region was further narrowed to a 31 Kb subunit of SD22-6, which contained palindromic AT-repeats (PATRRs). Interestingly, paralogs of SD22-6 flank copies of every other duplicon in the assembled alleles and each of these paralogs in hg38 contained PATRRs. Thus, we hypothesize that the PATRRs could be driving the rearrangements at this locus. PATRRs are known to form cruciform structures, which are prone to double strand breaks(*20*, *46*). If these breaks occur simultaneously at multiple loci in the genome, these are often resolved by non-homologous end joining (NHEJ), thereby rearranging the genome in some cases(*47*, *48*). PATRR is found to mediate constitutional chromosome 22 translocations, possibly via both replication dependent and independent mechanisms(*21*). Although replication independent mechanisms would imply that somatic mosaicism of PATRR mediated rearrangements would quickly arise, to date these have primarily been found to occur in sperm as a consequence of male meiosis(*49*, *50*). Only three cases for somatic mosaicism for 22q11 deletion have been reported (*51*, *52*). Similar to 22q translocations taking place in meiosis, LCR22 rearrangements could be the result of the same repair mechanism.

Previously, two models for chromosomal rearrangements were proposed: a breakage-first and a contact-first model(*53*). In the former, DNA breaks located at spatially long distances in the nucleus are moved towards each other to form a fusion chromosome. Such dynamic chromosomal movements have been shown to occur in an artificial translocation model for chromosome 22q11(*54*). Alternatively, first contact is made between (homologous) loci, which subsequently recombine. This is a likely scenario for the 22q11 deletion since aberrant pairing during meiosis followed by NAHR has been proposed as the deletion mechanism. The mechanism of recombination could involve the repair of double strand breaks caused by an unstable PATRR sequence. Moreover, if PATRRs are involved in the causal mechanism for the 22q11DS, this could explain the syndrome’s high frequency, since this is the only known locus genome-wide where numerous PATRRs and several stretches with high homology are localized in close vicinity(*55*).

While 22q11DS is the most frequent microdeletion syndrome, the underlying cause for the wide spectrum and variability of phenotypes observed has not been fully elucidated. Current approaches to reveal the cause of the phenotypic variability include targeted evaluations of single genes as well as genome-wide analyses in large cohorts of affected individuals(*56*). Variants in genes on the non-rearranged allele have been unmasked and might reveal autosomal recessive conditions(*57*). Variation of genes embedded in copy number variable regions like LCR22s has been so far ignored. We suggest that copy number variable genes embedded in the LCR22s could explain some of the phenotypic variability observed in individuals with the 22q11DS and human in general. Notably, SD22-3 contains at least two known active genes (TMEM191B and RIMBP3, **Fig. 1a**). TMEM191B is expressed in brain tissue (*58*, *59*). Not every allele of LCR22A features this duplicon, however neither is it observed to be present in more than one copy. Both TMEM191B and RIMBP3 have paralogs in LCR22D (TMEM191C, RIMBP3B and –C), but in the absence of an unambiguous reference sequence, it remains challenging to investigate the gene activity of each duplicon.

Other features embedded in the LCR22s may also influence the phenotypes of 22q11DS patients. The genes PRODH, DGCR5, and DGCR6 reside in SD22-5(*28*), which was retained in even the smallest mapped allele of LCR22A.. In all mapped individuals with the typical 22q11 deletion this duplicon is deleted, confirming previous observations of its hemizygosity in most patients(*28*, *29*, *60*–*62*). Additionally, the presence of pseudogenes (PI4KAP1 and P2, GGT3P, DGCR6L, BCRP2, GGT2, and lncRNAs (FAM230A, FAM230B and at least seven non-characterized paralogs) in different copy numbers could influence gene expression in a myriad of ways. Moreover, the observed size differences of LCR22A might exert a spatial effect on chromatin looping in the cell, thereby altering topologically associated domains(*63*–*65*). The phenotypic effect of variable repeat architecture could be minor for intact alleles, but could alter gene expression dramatically when the LCR22s are rearranged. The exact correlation between the LCR22 architectures and the genes embedded in them remains to be investigated. Hence, with these LCR22 assemblies, we envision future work to further elucidate the effect of multicopy genes at this locus.

In summary, optical mapping has allowed us to reveal an extraordinary level of variability within LCR22s. Our map of this genomic region is, to date, the most comprehensive for the LCR22s in the human genome reference sequence. Further, this map provides a framework for the alignment of both short and long read sequences which will ultimately close the remaining reference gaps and enable sequence-based analysis of the LCRs. Understanding the LCR variation will shed light on the mechanisms leading to 22q11 rearrangements and the different frequencies of the variation amongst populations. This knowledge will likely guide future prenatal counseling and testing for 22q11 related disorders. The LCR22s have rapidly expanded during hominoid evolution(*27*) and considering the region encompasses nine genes and 54 different RNAs, it seems plausible that the region influences important human traits. Thus, it is likely that the LCR variability has phenotypic consequences which may play a role in phenotypic variability in 22q11DS and affect other traits in the normal population. The ability to visualize and reconstruct complete and intact LCR haplotypes will greatly enhance our ability to start unraveling these important correlations.

## Material and Methods

### Patients & EBV cell lines

Patients with the 22q11DS were diagnosed using either a FISH assay with TUPLE1/ARSA probes (Abbot Molecular, Abbot Park, Illinois, USA), the MLPA SALSA P250 DiGeorge diagnostic probe kit (MRC-Holland), or with the CytoSure Constitutional v3 (4×180k) (OGT, Oxfordshire, UK). All individuals in the study were informed of the project’s outlines and gave written consent for their EBV cell lines and DNA to be used for sequencing and genotyping purposes. The study was approved by the Medical Ethics committee of the University hospital/KU Leuven (S52418), the Institutional Review Board approved research protocol (COMIRB # 07-0386) at the University of Colorado Denver, School of Medicine, and the Children’s Hospital of Philadelphia under Institutional Review Board (IRB) protocol 07-005352. Fiber FISH mapping was performed on Epstein Barr virus transformed lymphoblastoid cell lines from peripheral blood from probands and their parents. EBV cell lines were established as described^30^. Eleven patients were recruited during routine consultations in the hospital of Leuven, 1 at the Children’s Hospital of Philadelphia, and 2 at the Albert Einstein College of Medicine (IRB: 1999-201-047). HapMap control cell lines were obtained from the Coriell Cell Repository (Camden, NJ, USA) and cultured according to standard protocols

### Construction and screening of hybrid cell lines

Two individuals with the 22q11DS were randomly selected to produce stable, chromosome 22 haploid cell hybrids in a Wg3h Chinese hamster background. The latter is HPRT-deficient, allowing selection of fused cell colonies. Briefly, human EBV cells were fused with host cells in the presence of PEG and grown under selective HAT medium^31^. Cell colonies were screened by PCR and interphase FISH to select only those containing a del (22)(q11.21).

### *In Silico* Characterization of repeat subunits in LCR22s

All segmental duplication track positions were downloaded in bed format from UCSC in the region chr22:18,000,000-25,500,000 (hg38), including paralogous LCRs located elsewhere in the genome. These were merged with bedtools v2.17.0^32^ and sequences retrieved with the UCSC table browser^33^–^35^. These were then self-aligned using blastn v2.2.28+^36^ and filtered for reciprocal BLAST hits, alignments <100bp, and alignments <99% identity. If multiple queries aligned to the same subject segmental duplication at different positions the segmental duplication was split into multiple units. Unit positions were converted to bed format, sequences retrieved through the UCSC table browser, self-aligned again, and similarly filtered. Clusters of units aligning to each other were each considered a subunit family(**Table S1**).

The consensus of subunit families with at least one subunit between chr22:18,000,000-25,500,000 were created with Clustalx 2.1^37^ and the EMBOSS v6.5.7^38^ cons module. Bedtools was also used to subtract the subunits locations from the original UCSC region chr22:18,000,000-25,500,000 segmental duplication positions. The sequences of these “low percent identity” subunits were then retrieved using the UCSC table browser. The low percent identity sequences and subunit family consensus sequences were then merged into a final fasta file, repeat masked^39,40^, and sequences composed of >95% N’s removed.

### Extracting chr22 contigs from GM12878 10X *de novo* assembly data

*De novo* assembly data of BioProject ID PRJNA315896 were downloaded and searched for those contigs containing the first 50 Kb unique sequence flanking LCR22A, -B, -C, and –D. Next, contig fasta files were aligned to hg38 using BLAT^41^ and visualized in the UCSC genome browser to compare the transition of LCR22 repeats to unique flanking sequences to those in the reference genome.

### BAC DNA, Long range PCR Probe design and labeling

Using the subunit sequences library, fourteen fluorescent probes were designed (**Table S2**). Each probe targeted one subunit, which would also hybridize at the position of every paralogous subunit in that family. All those paralogous subunit positions were evaluated by BLAT alignment of probe sequences in the UCSC genome browser. Pattern colors were designed so that no two adjacent subunits in hg38 would produce the same fluorescent signal, in order to guarantee correct identification.

For each of these fourteen subunits long range PCR primer pairs were designed, producing amplicons between 2,946bp and 9,794bp (**Table S2**). PCR reactions were performed with the TAKARA LA v2 kit (Takara Bio inc.) using the standard gDNA protocol. Template gDNA was extracted from the same cell line for all reactions, in order to reduce amplicon variation between batches.

BAC clones were obtained from BacPac Resources (CHORI; Oakland) as *E.coli* stab cultures, which were grown according to recommendations. BAC DNA was extracted using the Nucleobond Xtra BAC kit (Macherey-Nagel). Subunit PCR Amplicons and BAC DNA were purified and antibody-labeled by random prime amplification (BioPrime DNA Labeling System; Invitrogen). An indirect detection system with primary labels Biotin-dUTP, Digoxigenin-dUTP, and Fluorescein-dUTP was used. The use of three labels allowed production of six detectable probe colors: three of each label separately and three of each pairwise combination.

### DNA combing, FISH, and fiber pattern assembly

DNA fibers were stretched using the GenomicVision extraction kit and combing system. Briefly, cultured EBV cells were embedded in an agarose plug and subsequently lysed and washed. Next, the agarose was dissolved and long DNA fragments were resuspended. Using an automated combing system, DNA fibers were consistently stretched on a coated glass coverslip by the receding meniscus method. YOYO-1 staining and scanning allowed visualization and evaluation of DNA fibers at this step.

Coverslips with combed DNA were hybridized with the designed probe pattern and washed using the manufacturer’s standard protocol. Probes were detected by indirect labeling with BV480 Streptavidin (pseudocolored red; BD Biosciences), Cy3 IgG Fraction Monoclonal Mouse Anti-Fluorescein (pseudocolored green; Jackson Immunoresearch), and Alexa Fluor 647 IgG Fraction Monoclonal Mouse Anti-Digoxigenin (pseudocolored blue; Jackson Immunoresearch). Probe mixes produced pseudocolors cyan, magenta, and yellow. Slides with labeled DNA were mounted in the provided scanner adapters and scanned at three excitation channels on a customized automated fluorescence microscope (GenomicVision).

Images were compiled to one complete slide recording and visualized in FiberStudio (GenomicVision). Slides were manually screened and fiber signals cropped to single image files. Individual images were visually aligned based on matching colors and distances between different probes. Fibers were tiled to complete alleles for LCR22A, B, C, and D, and compared to hg38 probe positions in the UCSC genome browser.

### Assembly of artificial LCR22 reference sequences

To confirm the fiber FISH assemblies, Bionano assays were performed on an overlapping cohort of 11 individuals. To compare results from the two methods, fiber FISH results were converted *in silico* into the optical map format. Using the hg38 reference genome sequences of SD22-1 to -5 and LCR22D, the sequence of each allele was predicted based on the orientation and copy number of subunits detected in the fiber FISH assemblies. Those sequences were then *in silico* labeled at recognition sites of the enzyme used for Bionano optical mapping, generating CMAP data files for all LCR22 repeats.

### High-molecular-weight DNA extraction

High molecular-weight DNA for genome mapping was obtained from whole blood. White blood cells were isolated from whole blood samples using a ficoll-paque plus (GE healthcare) gradient. The buffy coat layer was transferred to a new tube and washed twice with Hank’s balanced salt solution (HBSS, Gibco Life Technologies). A small aliquot was removed to obtain a cell count before the second wash. The remaining cells were resuspended in RPMI (Gibco Life Technologies) containing 10% fetal bovine serum (Sigma) and 10% DMSO (Sigma). Cells were embedded in thin low-melting-point agarose plugs (CHEF Genomic DNA Plug Kit, Bio-Rad). Subsequent handling of the DNA followed protocols from Bionano Genomics using the Bionano Prep Blood and Cell Culture DNA Isolation Kit. Briefly, the agarose plugs were incubated with proteinase K at 50°C overnight. The plugs were washed and then solubilized with GELase (Epicentre). The purified DNA was subjected to 45min of drop-dialysis, allowed to homogenize at room temperature overnight, and then quantified using a Qubit dsDNA BR Assay Kit (Molecular Probes/Life Technologies). DNA quality was assessed using pulsed-field gel electrophoresis.

### DNA labeling for Bionano genome mapping

The DNA was labeled using the Bionano Prep Early Access Direct Labeling and Staining (DLS) Kit (Bionano Genomics). DLS labels DNA using an epigenetic mark rather than by introducing single-strand nicks. Therefore, unlike the previously used enzyme Nt.BspQI, it does not create fragile sites that lead to consistent DNA breakage. 750 ng of purified genomic DNA was labeled by incubating with DL-Green dye and DLE-1 Enzyme in DLE-1 Buffer for 2 hr at 37°C, followed by heat-inactivation of the enzyme for 20 min at 70°C. The labeled DNA was treated with Proteinase K at 50°C for 1 hr, and excess DL-Green dye was removed by membrane adsorption. The DNA was stored at 4°C overnight to facilitate DNA homogenization and then quantified using a Qubit dsDNA HS Assay Kit (Molecular Probes/Life Technologies). The labeled DNA was stained with an intercalating dye and left to stand at room temperature for at least 2 hrs.

### Bionano data collection and assembly

The DNA was loaded onto the Bionano Genomics Saphyr Chip and linearized and visualized using the Saphyr system. The DNA backbone length and locations of fluorescent labels along each molecule were detected using Saphyr’s image detection software. Single-molecule maps were assembled *de novo* into genome maps using Bionano Solve with the default settings.

### Detection of structural variation within LCR22s

Structural variation in the LCR22s was evaluated in Bionano genome map data labeled using the Nt.BspQI nickase enzyme, from 154 individuals representing 26 diverse populations from five super-populations (Levy-Sakin et al. 2018). Assembled contigs mapping to LCR22s were realigned to chromosome 22 using RefAligner from BioNano Solve 3.1 with the following parameters: -res 2.9 -FP 0.6 -FN 0.06 -sf 0.20 –sd 0.0 -sr 0.01 -extend 1 -outlier 1e-14 -endoutlier 1e-13 -PVendoutlier -deltaX 12 -deltaY 12 -xmapchim 12 5000 -nosplit 0 -biaswt 0 -T 1e-12 -S -1000 -PVres 2 -rres 0.9 -MaxSE 0.5 -MinSF 0.15 -HSDrange 1.0 - outlierBC -outlierLambda 20.0 -outlierType1 0 -xmapUnique 12 -AlignRes 2. -outlierExtend 12 24 -Kmax 12 -resEstimate -M 1. The resulting alignments were visualized in OMView from the OMTools package42. The various haplotypes for the LCR22s were compiled and evaluated using the approach described in Supplementary Methods.

## Acknowledgements

Excellent technical assistance was provided by O.Tran and A. Jin. The molar cell line CHM1 was kindly provided by E. Eichler. **Funding:** W.D. is a fellow at KU Leuven supported by IWT (131625). This work was made possible by grants from the KUL PFV/10/016 SymBioSys and project 1665 Jerome Lejeune to J.R.V,. and GOA/12/015 to J.R.V. and K.D, by the National Institute of Mental Health (5U01MH101723-02), by the National Institute of General Medical Sciences (GM120772 to T.H.S and P.Y.K; GM125757 to B.S.E. and M.Z) and the Belgian Science Policy Office Interuniversity Attraction Poles (BELSPO-IAP) program through the project IAP P7/43-BeMGI. **Competing interests:** The authors declare to have no competing interests.

## Supplementary Materials

### Materials and Methods

Figures S1-S12

Tables S1-S2

## References

1. I. Human Genome Sequencing Consortium, Finishing the euchromatic sequence of the human genome. Nature. 431, 931–945 (2004).

2. J. A. Bailey et al., Recent segmental duplications in the human genome. Science (80-.). 297, 1003–1007 (2002).

3. J. A. Bailey, A. M. Yavor, H. F. Massa, B. J. Trask, E. E. Eichler, Segmental duplications: Organization and impact within the current human genome project assembly. Genome Res. 11, 1005–1017 (2001).

4. Z. Jiang et al., Ancestral reconstruction of segmental duplications reveals punctuated cores of human genome evolution. Nat. Genet. 39, 1361–1368 (2007).

5. M. Y. Dennis et al., The evolution and population diversity of human-specific segmental duplications. Nat. Ecol. Evol. 1, 0069 (2017).

6. M. Y. Dennis et al., Evolution of human-specific neural SRGAP2 genes by incomplete segmental duplication. Cell. 149, 912–922 (2012).

7. J. L. Boyd et al., Human-chimpanzee differences in a FZD8 enhancer alter cell-cycle dynamics in the developing neocortex. Curr. Biol. 25, 772–779 (2015).

8. C. Charrier et al., Inhibition of SRGAP2 function by its human-specific paralogs induces neoteny during spine maturation. Cell. 149, 923–935 (2012).

9. M. Florio et al., Human-specific gene ARHGAP11B promotes basal progenitor amplification and neocortex expansion. Science (80-.). 347, 1465–1470 (2015).

10. M. J. P. Chaisson et al., Resolving the complexity of the human genome using single-molecule sequencing. Nature. 517, 608–611 (2015).

11. D. Bovee et al., Closing gaps in the human genome with fosmid resources generated from multiple individuals. Nat. Genet. 40, 96–101 (2008).

12. G. Genovese et al., Using population admixture to help complete maps of the human genome. Nat. Genet. 45, 406–414 (2013).

13. K. Inoue, J. R. Lupski, Molecular Mechanisms for Genomic Disorders. Annu. Rev. Genomics Hum. Genet. 3, 199–242 (2002).

14. D. M. McDonald-McGinn et al., 22q11.2 deletion syndrome. Nat. Rev. Dis. Prim. 1, 15071 (2015).

15. L. Edelmann et al., A common molecular basis for rearrangement disorders on chromosome 22q11. Hum. Mol. Genet. 8, 1157–67 (1999).

16. C. Carlson et al., Molecular Analysis of Velo-Cardio-Facial Syndrome Patients with Psychiatric Disorders. Am. J. Hum. Genet. 60, 851–859 (1997).

17. T. H. Shaikh et al., Chromosome 22-specific low copy repeats and the 22q11.2 deletion syndrome: genomic organization and deletion endpoint analysis. Hum Mol Genet. 9, 489–501 (2000).

18. R. Weksberg et al., Molecular characterization of deletion breakpoints in adults with 22q11 deletion syndrome. Hum. Genet. 120, 837–45 (2007).

19. M. S. Hestand et al., A catalog of hemizygous variation in 127 22q11 deletion patients. Hum. Genome Var. 3, 15065 (2016).

20. D. R. F. Leach, Long DNA palindromes, cruciform structures, genetic instability and secondary structure repair. BioEssays. 16 (1994), pp. 893–900.

21. T. Kato, H. Kurahashi, B. S. Emanuel, Chromosomal translocations and palindromic AT-rich repeats. Curr. Opin. Genet. Dev. 22 (2012), pp. 221–228.

22. A. L. Gotter et al., A palindrome-driven complex rearrangement of 22q11.2 and 8q24.1 elucidated using novel technologies. Genome Res. 17, 470–481 (2007).

23. S. Lewis, E. Akgun, M. Jasin, Palindromic DNA and Genome Stability: Further Studiesa. Ann. N. Y. Acad. Sci. 870, 45–57 (1999).

24. L. A. Cunningham, A. G. Coté, C. Cam-Ozdemir, S. M. Lewis, Rapid, stabilizing palindrome rearrangements in somatic cells by the center-break mechanism. Mol. Cell. Biol. 23, 8740–50 (2003).

25. H. Kurahashi et al., Cruciform DNA structure underlies the etiology for palindrome-mediated human chromosomal translocations. J. Biol. Chem. 279, 35377–35383 (2004).

26. T. H. Shaikh et al., Low copy repeats mediate distal chromosome 22q11.2 deletions: Sequence analysis predicts breakpoint mechanisms. Genome Res. 17, 482–491 (2007).

27. X. Guo, L. Freyer, B. Morrow, D. Zheng, F. Index, Characterization of the past and current duplication activities in the human 22q11.2 region. BMC Genomics. 12, 71 (2011).

28. T. Guo et al., Deletion size analysis of 1680 22q11.2DS subjects identifies a new recombination hotspot on chromosome 22q11.2. Hum. Mol. Genet. 27, 1150–1163 (2018).

29. X. Guo et al., Variant discovery and breakpoint region prediction for studying the human 22q11.2 deletion using BAC clone and whole genome sequencing analysis. Hum. Mol. Genet., ddw221 (2016).

30. M. A. Eberle et al., A reference data set of 5.4 million phased human variants validated by genetic inheritance from sequencing a three-generation 17-member pedigree. Genome Res. 27, 157–164 (2017).

31. C. G. Cole et al., Finishing the finished human chromosome 22 sequence. Genome Biol. 9, R78 (2008).

32. V. A. Schneider et al., Evaluation of GRCh38 and de novo haploid genome assemblies demonstrates the enduring quality of the reference assembly. Genome Res. 27, 849–864 (2017).

33. L. Torres-Juan, J. Rosell, M. Sánchez-de-la-Torre, J. Fibla, D. Heine-Suñer, Analysis of meiotic recombination in 22q11.2, a region that frequently undergoes deletions and duplications. BMC Med. Genet. 8, 14 (2007).

34. K. A. Frazer et al., A second generation human haplotype map of over 3.1 million SNPs. Nature. 449, 851–861 (2007).

35. L. D. Botto et al., A population-based study of the 22q11.2 deletion: phenotype, incidence, and contribution to major birth defects in the population. Pediatrics. 112, 101–7 (2003).

36. D. M. McDonald-McGinn et al., The 22q11.2 deletion in African-American patients: An underdiagnosed population? Am. J. Med. Genet. 134 A, 242–246 (2005).

37. V. Goidts et al., Complex patterns of copy number variation at sites of segmental duplications: An important category of structural variation in the human genome. Hum. Genet. 120, 270–284 (2006).

38. R. E. Handsaker et al., Large multiallelic copy number variations in humans. Nat. Genet. 47, 296–303 (2015).

39. P. H. Sudmant et al., Global diversity, population stratification, and selection of human copy-number variation. Science (80-.). 349, aab3761 (2015).

40. M. Y. Dennis, E. E. Eichler, Human adaptation and evolution by segmental duplication. Curr. Opin. Genet. Dev. 41 (2016), pp. 44–52.

41. P. H. Sudmant et al., Diversity of human copy number variation and multicopy genes. Science. 330, 641–6 (2010).

42. K. M. Steinberg et al., Structural diversity and African origin of the 17q21.31 inversion polymorphism. Nat. Genet. 44, 872–80 (2012).

43. S. Aldridge et al., 1000 Genomes project. Nat. Biotechnol. 26, 256 (2008).

44. P. H. Sudmant et al., Diversity of human copy number variation and multicopy genes. Science (80-.). 330, 641–646 (2010).

45. V. Goidts et al., Identification of large-scale human-specific copy number differences by inter-species array comparative genomic hybridization. Hum. Genet. 119, 185–198 (2006).

46. K. S. Lobachev, D. A. Gordenin, M. A. Resnick, The Mre11 complex is required for repair of hairpin-capped double-strand breaks and prevention of chromosome rearrangements. Cell. 108, 183–193 (2002).

47. T. Kato et al., Two different forms of palindrome resolution in the human genome: Deletion or translocation. Hum. Mol. Genet. 17, 1184–1191 (2008).

48. H. Kurahashi, B. S. Emanuel, Long AT-rich palindromes and the constitutional t(11;22) breakpoint. Hum Mol Genet. 10, 2605–2617 (2001).

49. H. Kurahashi, B. S. Emanuel, Unexpectedly high rate of de novo constitutional t(11;22) translocations in sperm from normal males. Nat. Genet. 29, 139–140 (2001).

50. M. B. Sheridan et al., A palindrome-mediated recurrent translocation with 3:1 meiotic nondisjunction: The t(8;22)(q24.13;q11.21). Am. J. Hum. Genet. 87, 209–218 (2010).

51. A. Halder, M. Jain, M. Kabra, N. Gupta, Mosaic 22q11.2 microdeletion syndrome: diagnosis and clinical manifestations of two cases. Mol. Cytogenet. 1, 18 (2008).

52. M. A. Dempsey, S. Schwartz, D. J. Waggoner, Mosaicism del(22)(q11.2q11.2)/dup(22)(q11.2q11.2) in a patient with features of 22q11.2 deletion syndrome. Am. J. Med. Genet. A. 143A, 1082–6 (2007).

53. K. J. Meaburn, T. Misteli, E. Soutoglou, Spatial genome organization in the formation of chromosomal translocations. Semin. Cancer Biol. 17, 80–90 (2007).

54. V. Roukos et al., Spatial Dynamics of Chromosome Translocations in Living Cells. Science (80-.). 341, 660–664 (2013).

55. N. Delihas, A family of long intergenic non-coding RNA genes in human chromosomal region 22q11.2 carry a DNA translocation breakpoint/AT-rich sequence. PLoS One. 13, e0195702 (2018).

56. A. S. Bassett et al., Rare Genome-Wide Copy Number Variation and Expression of Schizophrenia in 22q11.2 Deletion Syndrome. Am. J. Psychiatry. 174, 1054–1063 (2017).

57. M. S. Hestand et al., A Catalog of Hemizygous Variation in 127 22q11 Deletion Patients. Hum. Genome Var. 3, 15065 (2016).

58. J. Lonsdale et al., The Genotype-Tissue Expression (GTEx) project. Nat. Genet. 45, 580–585 (2013).

59. L. Fagerberg et al., Analysis of the Human Tissue-specific Expression by Genome-wide Integration of Transcriptomics and Antibody-based Proteomics. Mol. Cell. Proteomics. 13, 397–406 (2014).

60. E. Michaelovsky et al., Genotype-phenotype correlation in 22q11.2 deletion syndrome. BMC Med. Genet. 13, 122 (2012).

61. H. Liu et al., Genetic variation at the 22q11 PRODH2/DGCR6 locus presents an unusual pattern and increases susceptibility to schizophrenia. Proc. Natl. Acad. Sci. U. S. A. 99, 3717–22 (2002).

62. H. Jacquet et al., The severe form of type I hyperprolinaemia results from homozygous inactivation of the PRODH gene. J Med Genet. 40 (2003) (available at http://www.jmedgenet.com/cgi/content/full/40/1/e7).

63. W. De Laat, D. Duboule, Topology of mammalian developmental enhancers and their regulatory landscapes. Nature. 502 (2013), pp. 499–506.

64. D. A. Kleinjan, V. van Heyningen, Long-Range Control of Gene Expression: Emerging Mechanisms and Disruption in Disease. Am. J. Hum. Genet. 76, 8–32 (2005).

65. J. Weischenfeldt, O. Symmons, F. Spitz, J. O. Korbel, Phenotypic impact of genomic structural variation: Insights from and for human disease. Nat. Rev. Genet. 14 (2013), pp. 125–138.

